# Accelerated Hematopoietic Stem Cell Aging in Space

**DOI:** 10.1101/2024.01.28.577076

**Authors:** Jessica Pham, Jane Isquith, Larisa Balaian, Luisa Ladel, Shuvro P. Nandi, Karla Mack, Inge van der Werf, Emma Klacking, Antonio Ruiz, David Mays, Paul Gamble, Shelby Giza, Jiya Janowitz, Trevor Nienaber, Tejaswini Mishra, Anna Kulidjian, Jana Stoudemire, Michael P. Snyder, Twyman Clements, Alysson R. Muotri, Sheldon R. Morris, Thomas Whisenant, Ludmil B. Alexandrov, Catriona H.M. Jamieson

## Abstract

Stem cell aging is accelerated by macroenvironmental and microenvironmental stressors, including inflammation. Previously, the NASA Twins study revealed inflammatory cytokine upregulation, chromosomal alterations, and telomere changes suggestive of accelerated aging in low-Earth orbit (LEO). To investigate the effects of spaceflight on human hematopoietic stem and progenitor cell (HSPC) aging, the NASA-supported Integrated Space Stem Cell Orbital Research team performed four independent 30- to 45-day NASA missions with matched flight and ground HSPC nanobioreactors in automated CubeLabs. These experiments revealed loss of HSPC dormancy, reduced self-renewal capacity, mitochondrial DNA amplification, APOBEC3-induced C-to-T mutagenesis, reduced ADAR1p150 expression, and alterations in the expression of repetitive elements. These molecular changes are indicative of accelerated HSPC aging and pre-leukemia stem cell generation in space and may be predictable and preventable.

## Introduction

Stem cells are functionally defined based on their capacity to self-renew, differentiate into tissue-specific progenitors, and become dormant in protective microenvironments or niches^1^. However, as the aging process unfolds, both the function and the quantity of hematopoietic stem cells (HSCs) change. HSC aging involves cell-intrinsic factors such as telomere attrition, acquisition of somatic mutations, changes in both epigenetic and epitranscriptomic (post-transcriptional) processes, and metabolic shifts, including changes in protein turnover regulation (proteostasis). These changes result in decreased stem cell regenerative capacity, their capacity to self-renew, and both innate and adaptive immune dysfunction characterized by a myeloid lineage bias^1-3^. Additionally, the acquisition of clonal mutations in HSCs, termed clonal hematopoiesis, contributes to immune dysfunction and pre-leukemia stem cell (pre-LSC) generation that ultimately fuels acute myeloid leukemia transformation and solid tumor progression^1, 4, 5^.

Cell-intrinsic changes that occur during HSC aging can occur following exposure to macroenvironmental stressors, such as environmental toxins and stress-inducing environments, inflammatory cytokine upregulation, and innate and adaptive immune disruption leading to chronic inflammation^1-3^. As tissue-specific stem cells and their progenitors are chronically exposed to inflammatory microenvironments, they become prone to hyperactivation of APOBEC3 base deaminases that introduce C-to-T mutations in the genome and ADAR1-mediated epitranscriptomic (post-transcriptional) A-to-I RNA editing^6^.

Additionally, seminal studies show that inflammatory microenvironments promote ADAR1-induced immune evasion and APOBEC3-mediated enzymatic mutagenesis in many human cancers^6-12^. To date, human whole genome, whole transcriptome, and single-cell sequencing analyses combined with stromal co-culture systems have revealed inflammatory cytokine-inducible APOBEC3 and ADAR1 as drivers of pre-cancer stem cell (CSC) and CSC generation^6, 10, 13-16^. Primate-specific APOBEC3 enzymes introduce C-to-T DNA base mutations in pre-LSCs in patients with MPNs^6, 11, 12^. Moreover, dual APOBEC3 and ADAR1 base deaminase deregulation has been linked to the oncogenic transformation of pre-LSCs into LSCs that harbor clonal self-renewal, survival, and cell cycle-altering genomic and epitranscriptomic alterations^17-21^. Also, a recent study showed that ADAR1 binds and edits R loops at telomere ends thereby contributing to genomic stability and longevity of malignant cells ^6, 22^. Together, stressful environmental conditions and advanced age have been associated with telomerase reverse transcriptase (TERT) activation, APOBEC3-induced DNA mutations, and ADAR1-mediated RNA modifications that promote pre-CSC and CSC longevity, malignant regeneration, and immune evasion by 20 different malignancies^6, 11, 13, 23-25^.

The National Aeronautics and Space Administration (NASA) Twins study revealed that a 340-day exposure to the macroenvironmental stress of low-Earth orbit (LEO) induced telomerase activation, pre-leukemic chromosomal translocations, and upregulation of inflammatory cytokines that are known to activate base deaminases^26^ capable of promoting pre-LSC and LSC generation^6, 13, 24^. Thus, protracted or frequent exposure to LEO may lead to persistent HSC dysfunction that triggers viral reactivation syndromes and pre-LSC as well as LSC generation. Accordingly, prediction of HSC fitness is vital for determining the risks of spaceflight and for developing countermeasures to prevent accelerated aging, latent viral reactivation, and cancer development in space. However, to date, few studies have investigated the effects of microgravity on biological processes regulated at the stem cell level^27^.

With NASA and implementation partner Space Tango, the Sanford Stem Cell Institute at the University of California San Diego established the first Integrated Space Stem Cell Orbital Research (ISSCOR) program to develop translational research platforms for studying hematopoietic, hepatic, and neural stem and progenitor cell physiology in microgravity and to develop countermeasures that curtail stem cell-related diseases. In this study, we report the development of a culture system enabling the long-term ex vivo expansion of human hematopoietic stem and progenitor cells (HSPCs) achieved through the utilization of a nanobioreactor. As of 2021, the ISSCOR program has successfully concluded four automated experiments using these biosensing human HSPC nanobioreactors that enabled monitoring stem cell fitness in LEO compared with ground controls with the aim of developing predictive biomarkers and countermeasures for accelerated and pre-malignant HSC aging.

## Results

### Development of Nanobioreactors for Hematopoietic Stem and Progenitor Cell Fitness Monitoring

While HSCs respond to injury and maintain the stem cell pool throughout life^28^, the NASA Twins study suggests that protracted exposure to microgravity accelerates tissue aging and induces immune dysfunction^29^. To study whether HSC fitness is impaired and immune dysfunction derives from the loss of normal stem cell homeostasis in space compared with terrestrial environments, we developed a biosensing 3D HSPC nanobioreactor system, enabling prolonged expansion of human HSPCs. To develop the biosensing 3D HSPC nanobioreactor system, we utilized fresh donor-derived HSPCs. Specifically, primary bone marrow samples were collected from individuals undergoing hip replacement surgery. Upon collection, mononuclear cells derived from the primary bone marrow were fractionated into CD34^-^ stromal and CD34^+^ HSPC-enriched fractions using magnetic bead selection. The CD34^-^ stromal fraction was irradiated and seeded in the nanobioreactor system that consists of a 3D SURGIFOAM® matrix in a gas-permeable pediatric blood bag. The CD34^+^ fraction, containing HSPCs, was cultured separately for 48 hours. To quantify cell cycle progression overtime at the single-cell level, CD34^+^ HSPCs were transduced with our lentiviral dual fluorescent ubiquitination-based cell cycle indicator 2 (FUCCI2BL), cell cycle sensing reporter^30^ directly following magnetic bead selection. Upon 48 hours, the HSPC-enriched fraction was seeded into nanobioreactors containing autologous stroma (**Fig. 1A**; see also **Fig. 1C**)^30^.

**Figure 1.**
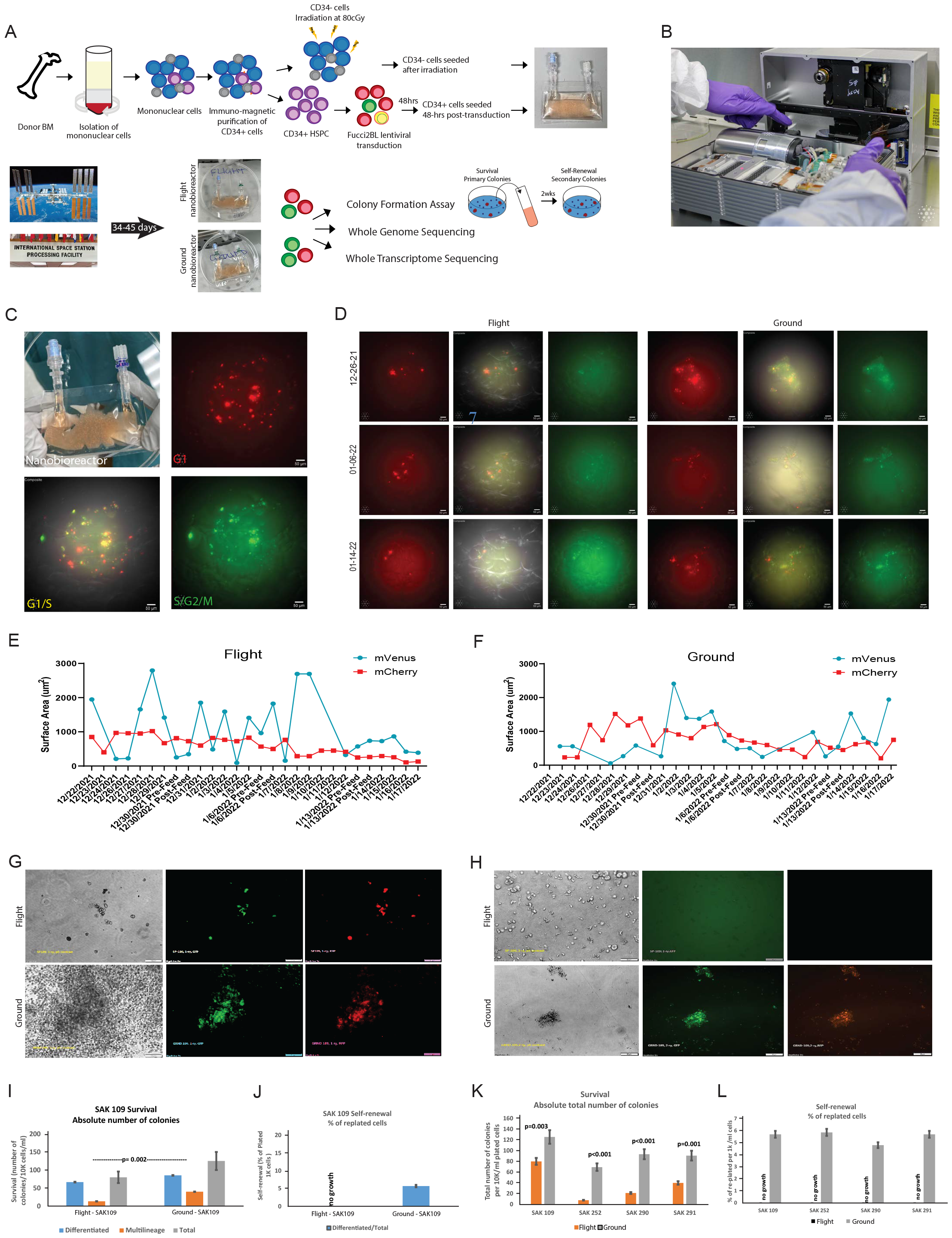
Paired nanobioreactors assess the effects of microgravity on hematopoietic stem and progenitor cells. **(A)** Experimental design of hematopoietic stem and progenitor cells (HSPCs) seeded in nanobioreactors for culture on SpaceX Commercial Resupply Service (SpX-CRS) missions to the International Space Station (ISS). (B) The CubeLab integrated system is equipped with a thermal management system, a microscope imaging system known as TangoScope, and fluid routing systems to transport media to and from cells. (C) FUCCI2BL transduced HSPCs imaged within the nanobioreactor. mVenus-hGem indicates S/G2/M phase and mCherry-hCdt1 indicates G1 phase of the cell cycle. mVenus+/mCherry+ cells indicate G1/S phase. (D) Representative fluorescent images acquired <u>during</u> SpX-CRS24 mission. Images were taken at the same X-Y-Z position within each nanobioreactor to capture the cell cycle transit of aged bone marrow (ABM) CD34^+^ HSPCs. Images were taken at 20X. (E-F) Representative Volocity fluorescent intensity quantification measured in paired SAK-109 flight and ground nanobioreactors <u>over the course of</u> SpX-CRS24 mission. Fluorescence was quantified in mCherry- and mVenus-expressing CD34^+^ HSPCs and measured as total surface area (µm^2^). (G-H) Colony-formation and replating assay of flight and ground returned samples <u>after</u> SpX-CRS24 mission. (G) Representative images of primary colonies-HSPC survival assay (H) Representative images of secondary colonies-HSPC self-renewal assay. (I-J) Representative ABM derived mononuclear cells (SAK-109) were cultured in parallel in flight and ground nanobioreactors for 42 days, then subjected to clonogenic assays, as described in Fig 1a and Methods. (I) In survival assay (primary colonies) the graph demonstrates the absolute number of differentiated and multilineage colonies, which were counted separately. (J) In self-renewal assay (secondary colonies) the graph depicts the % of replated cells. Each bar represents Mean+/-SE for triplicate conditions. Statistical analysis included Student’s t-test and one-way ANOVA, including All Pairwise Multiple Comparison Procedures (Holm-Sidak method). (K-L) Summarized bar graph of four individual paired ABM samples (SAK-109, SAK-252, SAK-290, and SAK-291) cultured in parallel in flight and ground nanobioreactors for 42 days, then subjected to clonogenic assays, as described in Fig 1a and Methods. (K) In survival assay the graph demonstrates the absolute total number of colonies. (L) In Self-renewal assay the graphs depict the % of replated cells. Each bar represents Mean+/-SE for triplicate conditions. Statistical analysis included Student’s t-test and one-way ANOVA, including All Pairwise Multiple Comparison Procedures (Holm-Sidak method).

To enable continuous confocal fluorescence microscopic imaging and ensure the viability of HSPCs in nanobioreactors, we utilized Space Tango CubeLab™ hardware that incorporates Takasago peristaltic pumps to simulate physiologically relevant flow at 2 mL/min during the once-weekly media exchanges (**Fig. 1B, S1A**)^31^. The CubeLab included a TangoScope™ confocal fluorescence microscope with a 20X objective for dual-color fluorescence imaging to assess cell cycle alterations with the FUCCI2BL reporter and sensors to monitor CO_2_ levels and 37°C temperature to ensure consistent (nominal) culture conditions (**Fig. S1B**). To assess changes associated with microgravity, nanobioreactors were seeded and integrated into paired CubeLabs that were destined for flight on NASA Commercial Resupply Service (CRS) missions by SpaceX or maintained at the Kennedy Space Center’s Space Station Processing Facility under the same 37°C and 5% CO_2_ culture conditions. In four independent NASA SpaceX CRS (SpX-24, SpX-25, SpX-26, and SpX-27) missions, ranging in duration from 32 to 45 days, nanobioreactors containing HSPCs and stromal cells from paired and unpaired samples were flown and, where applicable, compared with terrestrial nanobioreactor controls (**Table 1**).

**Table 1.** NASA ISSCOR Aged Normal Bone Marrow Patient Characteristics

| Mission | Launch Date | Splashdown Date | Mission Duration | Post-Processing Date | Total Experiment Duration | Sample | Condition | Age | Sex |
| --- | --- | --- | --- | --- | --- | --- | --- | --- | --- |
| SpX-CRS24 | 12/21/2021 | 1/24/2022 | 34 days | 1/26/2022 | 42 days | SAK-109 | *Paired Flight & Ground* | 76 | F |
| SpX-CRS24 | 12/21/2021 | 1/24/2022 | 34 days | 1/26/2022 | 42 days | SAK-072 | Ground | 65 | F |
| SpX-CRS24 | 12/21/2021 | 1/24/2022 | 34 days | 1/26/2022 | 42 days | SAK-066 | Flight | 75 | F |
| SpX-CRS25 | 7/14/2022 | 8/20/2022 | 37 days | 8/24/2022 | 50 days | SAK-201 | Paired Flight & Ground | 67 | F |
| SpX-CRS25 | 7/14/2022 | 8/20/2022 | 37 days | 8/24/2022 | 50 days | SAK-212 | Paired Flight & Ground | 55 | F |
| SpX-CRS26 | 11/26/2022 | 1/11/2023 | 45 days | 1/13/2023 | 60 days | SAK-252 | Paired Flight & Ground | 60 | M |
| SpX-CRS27 | 3/14/2023 | 4/15/2023 | 32 days | 1/13/2023 | 42 days | SAK-290 | Paired Flight & Ground | 74 | F |
| SpX-CRS27 | 3/14/2023 | 4/15/2023 | 32 days | 4/18/2023 | 42 days | SAK-291 | Paired Flight & Ground | 91 | M |

### Accelerated Hematopoietic Stem Cell and Progenitor Cycle Kinetics in Space

To study single stem cell cycle kinetics in LEO compared with terrestrial ground controls, CD34^+^ cells were lentivirally transduced with the FUCCI2BL reporter (**Fig. 1C**)^30^. Cell cycle transit times were analyzed by daily confocal fluorescence microscopic imaging of FUCCI2BL reporter activity in flight and ground nanobioreactors in the same pre-set locations within the nanobioreactors using a three-axis microscope system (TAMS) and artificial intelligence (AI) tracking with a GPU-accelerated embedded processor in the CubeLab (**Fig. S1C, video**). Volocity software-based quantification revealed that HSPCs in space exhibited decreased cell cycle transit times compared to their paired ground controls. This was evident through dynamic alterations in the mVenus signal, signifying proliferating cells in G1/S phase (Fig. 1D-1F). Additionally, HSPCs in space exhibited reduced mCherry expression, indicating cells in G1 phase and a loss of quiescence, characteristic of accelerated HSC aging.

### Space Impairs Hematopoietic Stem and Progenitor Cell Function

Prior to each mission, CD34^+^ HSPCs from the pool of mission-ready samples were utilized for *in vitro* survival and replating assays to establish baseline levels of clonogenicity and self-renewal capacity (**Fig. S1D-E**). Upon completion of the mission, nanobioreactors were de-integrated from the flight and ground CubeLabs and returned within 48 to 96 hours to the UC San Diego ISSCOR laboratory under consistent (nominal) culture conditions for processing (**Fig. S1F**). Cells were flushed from the nanobioreactors and utilized for colony forming and replating assays (**Fig. 1G-H**). Compared with ground controls, samples exposed to LEO had both reduced colony survival and replating capacity indicative of accelerated HSC aging in space (**Fig. 1I-L, S1G-H**). Additionally, functional, genomic, and transcriptomic analyses were performed on cells returning from one of four SpX-CRS missions, denoted as post-flight samples.

### Whole Genome Sequencing Detection of Space-Associated Telomere Attrition

To gain further insights into the accelerated aging of HSCs in microgravity, cells from flight and ground nanobioreactors were sent for whole genome sequencing with 150 bp pair-end reads and 90X coverage to detect the accumulation of somatic mutations and assess changes in telomere length (**Fig. S2A**). As cells age, telomeres shorten with every cell division^32^. Previous studies have shown that regardless of time spent in LEO, telomeres shorten upon return to Earth^33^. By utilizing a robust computational approach to measure telomere length, TelomereCat (**Fig. 2A**), we observed telomere attrition in post-flight samples compared with matched terrestrial controls (**Fig. 2B-C**). To corroborate this trend, we performed whole transcriptome sequencing (RNA-seq) with Gene Set Enrichment Analysis (GSEA) to identify telomere-associated pathway genes that were differentially expressed (**Fig. 2D, S2B**). Through this analysis, we identified six significantly differentially expressed genes (q-values<0.05) related to the telomere pathway, four of which were downregulated post-flight and two of which were upregulated post-flight. Overall, these results indicated the potential for telomere attrition and accelerated aging in space.

**Figure 2.**
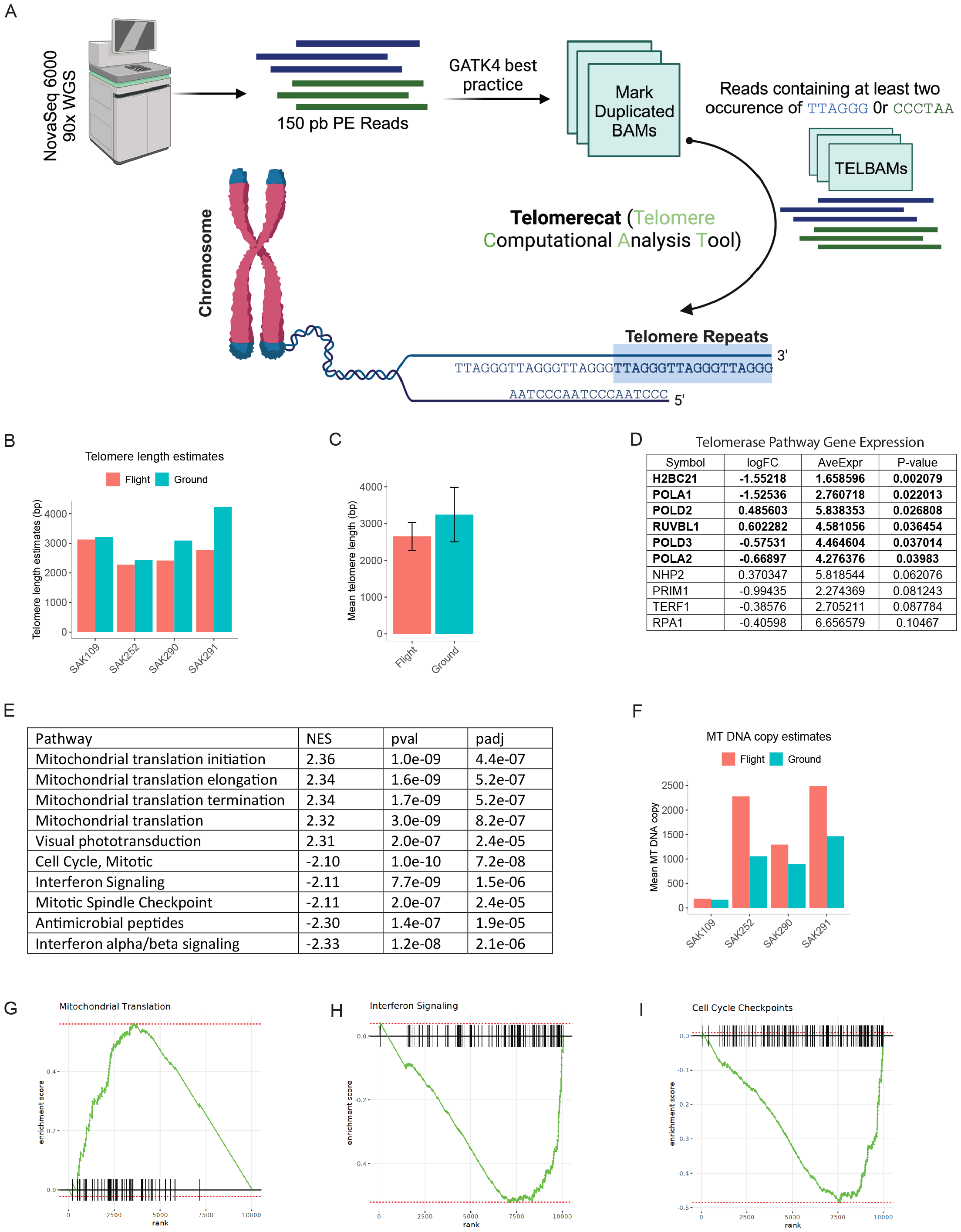
Genomic stressors linked to microgravity. **(A)** Schematic of telomere length estimation from whole genome sequencing data. Raw FASTQ files were downloaded within Triton shared computational cluster environment. Next, the GRCh38.d1.vd1 reference sequence was used for the alignment of raw reads. Subsequently, following GATK4 best practice, Telomerecat was employed for estimation of telomere length. (B) Bar graphs illustrating the Telomerecat-estimated overall telomere length (in base pairs (bp)) for hematopoietic stem and progenitor cells (HSPCs). These cells are derived from paired flight and ground samples of SAK-109, SAK-252, SAK-290, and SAK-291 upon their return. (C) Bar plots of mean telomere length estimates (in bp) from the four replicates shown in Figure 2B. Bar plots represent the mean ± SEM. (D) Summary of top expression differences (measured by limma p-value) for telomere associated pathway genes curated from the Reactome pathways HSA-157579, HSA-171319, and HSA-171306 (E) Gene set enrichment analysis (GSEA) results for comparison of flight (SAK-066, SAK-109, SAK-252) and ground (SAK-072, SAK-109, SAK-252) RNA-Seq samples using Reactome pathways as ontological database. NES - Normalized Enrichment Score (F) Bar plots showing mitochondrial DNA copy number average for the flight vs ground samples from HSPC (hematopoietic stem progenitor cells) samples produced by fastMitoCalc. (G-I) GSEA enrichment plots for the comparison of ground and flight RNA-Seq samples show the distribution of the enrichment score (ES, green line) and the position of each gene in the set across the gene ranks for the Mitochondrial Translation (G), Interferon Signaling (H), and Cell Cycle Checkpoints (I) pathways. The Mitochondrial Translation pathway has a positive normalized ES (NES) while the other two pathways have negative NESs.

### Mitochondrial Stress Responses Increase in Space

Considering that mitochondrial stress responses are indicative of HSC aging, we performed whole transcriptome (RNA-seq) analyses with Reactome-pathway-based GSEA that revealed significantly increased mitochondrial-related gene expression in post-flight compared with ground control nanobioreactor samples (**Fig. 2E**). To corroborate these findings, we performed whole genome sequencing (WGS) analyses at 90X coverage to quantify mitochondrial DNA (mtDNA) copy number (**Fig. 2F-2G, S2C**). Notably, HSPCs cultured in LEO harbored mtDNA copy number amplification compared to ground controls. Detection of mtDNA copy number amplification combined with a reduction in cell cycle checkpoint expression was indicative of stress-induced proliferative HSPC responses in microgravity that could reduce HSC fitness and accelerate aging (**Fig. 2H-2I, S2D-S2E**).

### Space Activates APOBEC3-mediated C-to-T DNA Mutagenesis

Somatic mutations in microgravity were identified using WGS analyses of cells obtained from paired post-flight and ground samples (n=4 pairs). Single base C-to-T substitutions (SBSs) represented the most common mutations (**Figs. 3A-E, Fig. S3A**). Insertions and deletions (indels) were infrequent and only one sample (SAK109) showed doublet base substitutions (DBSs) (**Figs. 3A-E, Fig. S3A**). To determine if microgravity-associated C-to-T mutagenesis conformed with an APOBE3C mutagenesis pattern, CD34^+^ cells from three aged bone marrow samples were lentivirally transduced with APOBEC3C or pCDH lentiviral backbone controls ^34^. Notably, APOBEC3C overexpression was primarily comprised of SBSs and mirrored the mutation patterns seen in flight samples (**Fig. 3F-J and fig. S3B**). These patterns were also similar to APOBEC3C mutagenesis patterns detected in MPN pre-LSC populations thereby suggesting that microgravity may induce pre-leukemic mutations by activation of stress enzymes^6^. Somatically and non-synonymously mutated genes were identified within HSPC and APOBEC3C cohorts (**fig. S3C-F)**. Overlapping mutated genes between HSPC nanobioreactors post-flight and lentiviral APOBEC3C transduced CD34^+^ cells were identified; notably, the top four differentially mutated genes (DPP6, IL1RAPL1, NYAP2, and PDE4D) have distinctive roles in neuronal function and stress response genes involved in tissue homeostasis (**fig. S3G-H)**.

**Figure 3.**
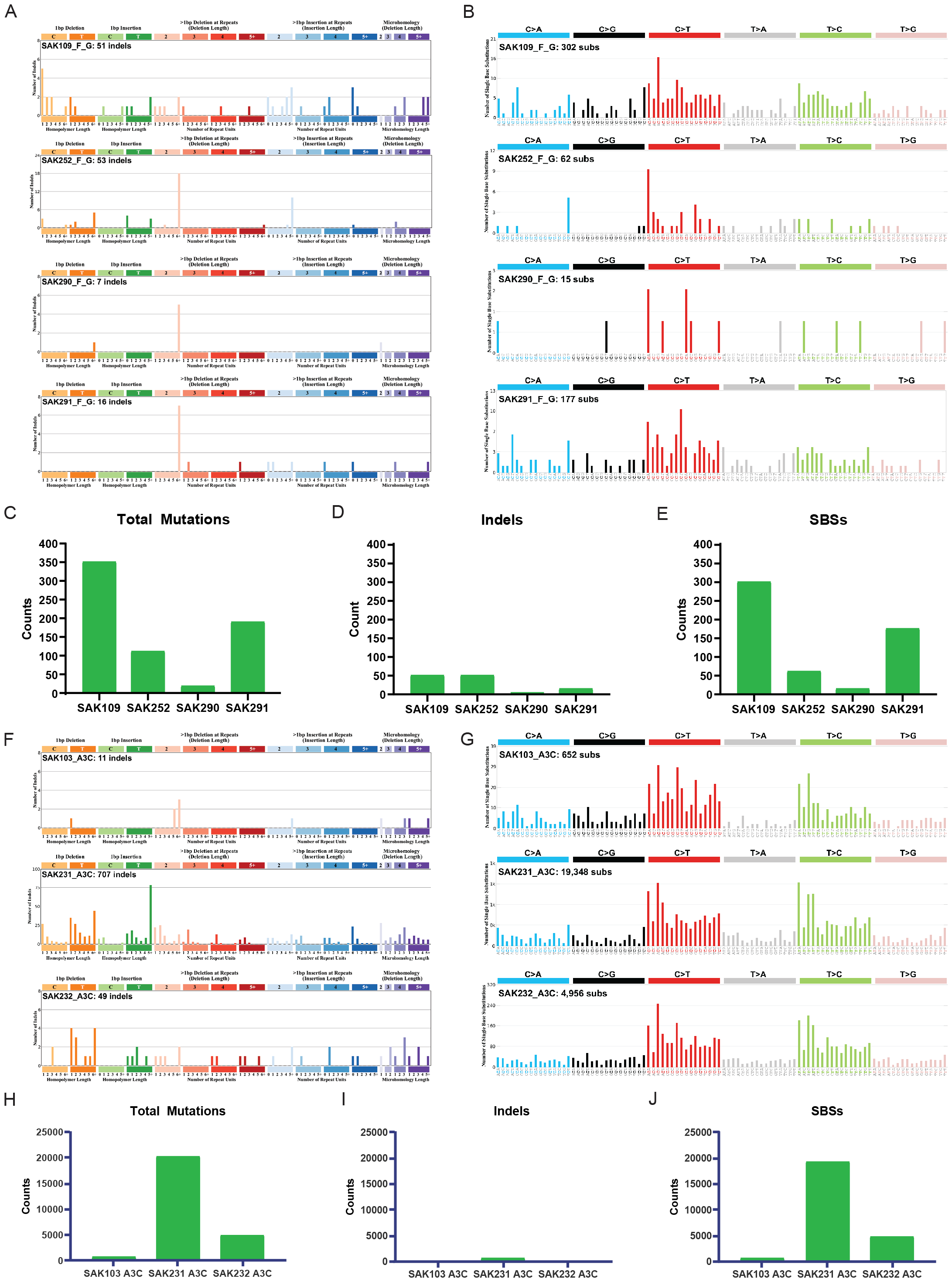
Mutation patterns acquired in hematopoietic stem and progenitor cells after 30-days in low-Earth orbit. **(A)** Patterns of small insertions and deletions (indels) from hematopoietic stem progenitor cells (HSPCs) are shown using the ID-83 classification scheme on the x-axis. The y-axis is scaled differently in each plot to optimally show each mutational pattern with the y-axis reflecting the number of mutations for the respective mutational scheme. The data from returned flight samples is normalized to returned ground data. (B) Patterns of single base substitution (SBS) for the samples from HSPCs are shown using the SBS96 classification scheme on the x-axis. The y-axis is scaled differently in each plot to optimally show each mutational pattern with the y-axis reflecting the number of mutations for the respective mutational scheme. The data from returned flight samples is normalized to returned ground data. (C) Bar plots showing the total amount of somatic mutations acquired in flight in comparison to its ground control. The Y-axis reflects the amounts of total mutation measured in somatic mutation counts. The X-axis corresponds to the four replicated experiments. (D) Bar plots showing the total amount of somatic insertion and deletion (indel) mutations acquired in flight in compared to its ground control. The Y-axis reflects the amounts of total small insertions and deletions (indels), measured in somatic mutation counts. The X-axis corresponds to the four replicated experiments. (E) Bar plots showing the total amount of somatic single base substitution (SBS) mutations acquired in flight in comparison to its ground control. Y-axis reflects the amounts of total SBS, measured in somatic mutation counts. X-axis corresponds to the four replicated experiments. (F) Patterns of small insertions and deletions (indels) for samples from APOBEC3C containing pCDH vector transduced cells are shown using the ID-83 classification scheme on the x-axis. The Y-axis is scaled differently in each plot to optimally show each mutational pattern with the y-axis reflecting the number of mutations for the respective mutational scheme. (G) Patterns of single base substitution (SBS) for the samples from APOBEC3C containing pCDH vector transduced HSPCs are shown using the SBS96 classification scheme on the x-axes. Y-axes are scaled differently in each plot to optimally show each mutational pattern with the y-axes reflecting the number of mutations for the respective mutational scheme. (H) Bar plots showing total amount of somatic mutations acquired in APOBEC3C containing pCDH vector transduced cells. The Y-axis reflects the amounts of total mutation measured in somatic mutation counts. The X-axis corresponds to the four replicated experiments. (I) Bar plots showing the total amount of somatic indel mutations acquired in APOBEC3C containing pCDH vector transduced cells. The Y-axis reflects the amounts of total small insertions and deletions (indels), measured in somatic mutation counts. The X-axis corresponds to the four replicated experiments. (J) Bar plots showing the total amount of somatic single base substitution (SBS) mutations acquired in APOBEC3C containing pCDH vector transduced cells. The Y-axis reflects the amount of total SBS, measured in somatic mutation counts. The X-axis corresponds to the four replicated experiments.

### Base Deaminase and Repetitive Element Expression Changes in Space

Additionally, we performed RNA-seq analyses using established bioinformatics pipelines^10, 24^. Overall, we observed a decrease in gene expression in post-flight HSPC nanobioreactors compared with their paired ground controls (**Fig. 4A**). Subsequent, splicing analysis showed that exon skipping and intron retention were approximately the same in post-flight compared with ground samples (**Fig. 4B**). However, ADAR1 p150/ADAR1 p110 splice isoform ratios were significantly decreased in post-flight samples (p-value=0.036) (**Fig. 4C**). The reduction in ADAR1p150 to p110 ratios in post-flight compared to ground samples corresponded with decreased A-to-I RNA edits per million (p-value=0.007) (**Fig. 4D**) and correlated with a reduction in replating capacity (**Fig. 1G-H)** in keeping with the role of ADAR1p150 in HSPC self-renewal regulation^10^. Moreover, RNA-seq analysis revealed upregulation of APOBEC3A and APOBEC3C in post-flight compared to ground control samples (**Fig. 4E, Fig. S4A; Fig. S4B**). This result was consistent with RNA-seq data from the NASA Twins study^26^, which showed that APOBEC3A was up-regulated in lymphocyte-depleted cells as time in space increased (**Fig. S4F**). Notably, ADAR1 was down-regulated in lymphocyte-depleted cells as time in microgravity increased (**Fig. S4G**) suggesting that base deaminase deregulation may promote accelerated aging and pre-malignant mutations in LEO.

**Figure 4.**
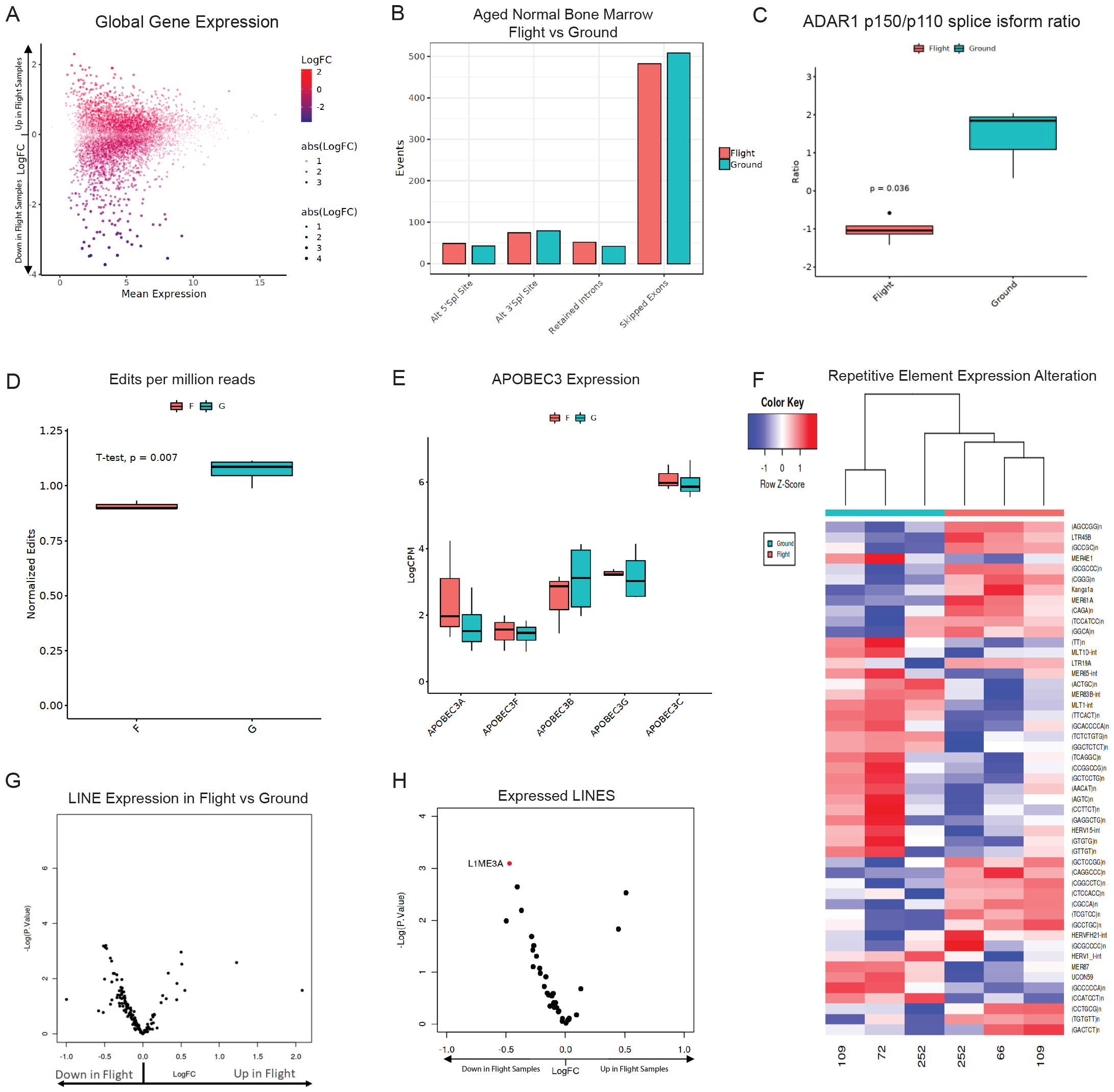
Gene expression changes resulting from microgravity. **(A)** Plot visualizing mean expression (log counts per million (lcpm)) and log fold change (LogFC) for comparison of hematopoietic stem and progenitor cells (HSPCs) derived from flight and ground samples upon return. (B) Summary of significant differential splicing events (FDR < 0.05, |PSI| > 0.1) for the comparison of HSPCs derived flight and ground samples upon return. (C) Expression of the ratio of ADAR p150 and ADAR p110 lcpm values in HSPCs derived flight and ground samples upon return. Student’s t-test was performed to determine significance. (D) Summary of detected RNA editing events normalized by library size and sequencing run for HSPCs derived flight and ground samples upon return. Student’s t-test was performed to determine significance. (E) Expression of detectable APOBEC3 genes measured by lcpm in HSPCs derived flight and ground samples upon return. (F) Heatmap of repetitive elements with p-value < 0.01 in HSPCs derived flight and ground samples upon return. (G) Volcano plot of HSPCs derived flight and ground samples upon return visualizing LINE expression levels. The Y-axis represents the negative log of the p-value (-Log P Value) and the x-axis is the log fold change (LogFC). (H) Volcano plot of LINE element expression in Flight vs Ground RNA-Seq samples for LINEs that are differentially expressed in the APOBEC3C overexpression vs pCDH comparison. The Y-axis represents the negative log of the p-value (-Log P Value) and the x-axis is the log fold change (LogFC). Red dots represent repetitive elements with p-value < 0.05.

Deregulation of APOBE3 and ADAR1 base deaminases can alter repetitive element expression and has been linked to accelerated aging^35^. To elucidate if repetitive element expression changes occur in space, we performed differential expression analysis of repetitive elements specifically. This analysis revealed decreased LINE element expression between post-flight and ground samples (**Fig. 4F-G**). In keeping with LINE element restriction by APOBEC3 enzymes, lentiviral APOBEC3C transduced CD34^+^ cells differentially expressed LINEs compared with backbone transduced CD34^+^ cells (**Fig. 4H, Fig. S4E**). Of the 38 differentially expressed LINEs within the APOBEC3C WT compared with pCDH transduced control samples, L1ME3A was significantly repressed (p-value<0.05) in post-flight samples compared to ground controls (**Fig. 4H**).

## Discussion

Few studies have examined the effects of microgravity on human stem cell-specific biological processes^27^. In this study, we report the development of a culture system enabling the long-term ex vivo expansion of human hematopoietic stem and progenitor cells (HSPCS) achieved through the utilization of a nanobioreactor. Results from our study suggest that genomic, epi-transcriptomic (post-transcriptional), and mitochondrial changes may cause loss of HSPC dormancy, reduced self-renewal capacity, mitochondrial DNA amplification, APOBEC3-induced C-to-T mutagenesis, reduced ADAR1p150 expression, and alterations in expression of repetitive elements, which taken together, may contribute to accelerated HSPC aging. Microgravity activates stress response genes, APOBEC3C and ADAR1, which mutate DNA and alter RNA. ADAR1 splice isoform switching to the malignant, cytokine-inducible, A-to-I editing, ADAR1p150 isoform is implicated in disease progression such as myeloproliferative neoplasms^16^. During normal hematopoietic tissue homeostasis and multi-lineage reconstitution, the potential of HSCs is regulated in part by ADAR1 activity^36^. The decrease in ADAR1p150 activity may be linked to the significantly reduced self-renewal capacity of post-flight samples. Indeed, decreased A-to-I editing corroborated the significant decrease in expression of ADAR1p150 activity. These findings are consistent with what was observed in the NASA Twins Study transcriptomics data showing ADAR1 in lymphocyte-depleted cells was down-regulated as time in microgravity increased^26^.

In extensive whole genome sequencing analyses, we observed an overlap in synonymous and non-synonymous mutations between spaceflight and lentiviral APOBEC3C-transduced CD34^+^ cells. These mutations centered primarily within neuronal regulatory genes, including DPP6, IL1RAPL1, NYAP2, and PDE4D)^37-40^. Recalling the observed contribution of nociceptive neurons and sympathetic innovation of the bone marrow niche to stress responses, it may be reasonable to suggest that the neuronal genes seen in our study potentially play a role in regulating HSC maintenance, in particular the “fight or flight” response seen in bone marrow under conditions of extreme stress^41^. Functional changes in HSCs accelerate aging with myeloid lineage skewing and decreased innate and adaptive immune responsiveness, setting the stage for the loss of cancer immune surveillance.

The mutations acquired in post-flight samples may also result from APOBEC activation, giving significance to the LINE-1 element expression seen in the APOBEC3C WT overexpressing cells. Human LINE-1 elements are seen in many cancers and the most active L1 elements have been shown to evade somatic repression^42^. APOBEC3 has been shown to repress LINEs and restrict replication^43^; L1ME3A is differentially regulated in young versus senescent human umbilical vein endothelial cells (HUVECs)^44^ and was repressed in post-flight samples. Genetic and epigenetic changes and immune responses elicited by retrotransposon reactivation have been linked to aging^35^ and cancer^44^. Data from the NASA Twins study showed that APOBEC3A was up-regulated in lymphocyte-depleted cells as time in space increased^26^. Mitochondrial stressors have also been seen in several astronaut health studies in the context of peripheral blood^33, 45, 46^. The returning flight samples notably had acquired C-to-T single-base substitution mutations previously linked to APOBEC activation^34^. Strikingly, these mutations share similarities to mutation profiles of pre-leukemic stem cells^6^.

While the etiology of pre-leukemic changes in the blood was not clearly elucidated in the NASA Twins study^26^ our research suggests that LEO is a uniquely accelerating environment for understanding stem cell dysfunction that leads to pre-cancer initiation and loss of immune surveillance against cancer. Further, we found that the effects of microgravity on astronauts can be recapitulated *in vitro* at the stem cell level, providing the impetus for utilizing space to dissect mechanisms of immune dysfunction and pre-LSC generation in a compressed timeframe^47^. We are currently studying the potential to identify viable regenerative elements in aged normal bone marrow stem cells post-flight, which would indicate that the microgravity environment does not necessarily suppress cell function; however, further analysis is needed to determine this relationship and that of any observed inflammatory responses and immune system activation.

Microgravity can also accelerate medical advances on Earth such as drug development, manufacturing of biomedical products, and cellular therapies. For future missions, it would be interesting to monitor molecular changes in real-time by using fluorescent-based reporters in different tissues and cellular contexts. The International Space Station (ISS) environment provides a unique opportunity to study stem cell aging and the development of immune dysfunction, which will enhance the safety of space exploration and for pre-CSC detection and eradication terrestrially. Since 600 astronauts have flown to the ISS and commercial LEO platforms are becoming more accessible, the development of predictive biomarkers and countermeasures for accelerated aging and pre-cancer development will have significant value in space and terrestrially.

(I)Bar plots showing the total amount of somatic indel mutations acquired in APOBEC3C containing pCDH vector transduced cells. The Y-axis reflects the amounts of total small insertions and deletions (indels), measured in somatic mutation counts. The X-axis corresponds to the four replicated experiments.

(J)Bar plots showing the total amount of somatic single base substitution (SBS) mutations acquired in APOBEC3C containing pCDH vector transduced cells. The Y-axis reflects the amount of total SBS, measured in somatic mutation counts. The X-axis corresponds to the four replicated experiments.

## Methods

### Nanobioreactor

Nanobioreactors were constructed from 2-port, fluoroethyl polymer (FEP) film bags that are gas permeable, chemically inert, and transparent for imaging (OriGen Biomedical). A porcine gelatin SURGIFOAM (J&J MedTech) matrix was cut to size to function as a physical scaffold (trabecular bone) before resealing with a heat sealer under sterile conditions.

### Primary Patient Sample Preparation for Seeding Nanobioreactors

Aged normal bone marrow samples were collected from routine hip replacement surgeries under an IRB-approved protocol. Mononuclear cells were isolated by Ficoll-Pacque density centrifugation and either cryopreserved in liquid nitrogen or immediately followed by CD34^+^ selection. CD34^+^ cells were selected using MACS magnetic bead separation according to the manufacturer’s protocol (Miltenyi, catalog 130-100-453). The CD34^-^ fraction was resuspended in HBSS + 2% FBS prior to irradiation at 80cGy. The irradiated negative fraction was passed through a 70uM filter to create a single-cell suspension prior to counting and seeding at 2e6 cells per nanobioreactor in StemPro media (Stem Cell Technologies Inc). The CD34^+^ fraction was resuspended in 1X StemPro-34 SFM medium supplemented with a cytokine cocktail of SCF (50ng/ml), TPO (10ng/ml), FLT3 (50ng/ml), and IL-6 (10ng/ml). CD34^+^ cells were plated at 55-70K cells per 96-well U-bottom plate and transduced with Fucci2BL lentivirus at a MOI of 50-80. After 48-hours, 600K CD34^+^ Fucci2BL transduced cells were seeded into nanobioreactors pre-cultured with the allogenic CD34^-^ cell fraction and StemPro media (Stem Cell Technologies Inc.).

### Shipping Samples to Kennedy Space Center

Fully seeded nanobioreactors were shipped from San Diego, California to Kennedy Space Center via FedEx First Overnight. Samples were loaded into a MicroQ iQ2 unit supporting 37°C temperature control (MicroQ Technologies). Humidity was maintained by dampening Kimwipes with sterile water and padding the unit chamber. Once the nanobioreactors were placed into the chamber, the chamber was purged with 5% CO_2_ before sealing for shipment.

### CubeLab operations (flight and ground, all missions)

Space Tango flight and ground hardware support two nanobioreactors and automated operations (fluid conveyance, fluid routing, thermal control, environmental monitoring, imaging, and sampling) within each 9U CubeLab™. All wetted materials selected for use within the CubeLab were certified biocompatible and sterilizable. Wetted materials that come in contact with fluids or cells were sterilized, assembled, and maintained sterile, tested, and confirmed to be free of microbial contamination prior to loading experimental fluids and biology.

Science and CubeLabs (flight and ground) were prepared at the Kennedy Space Center’s Space Station Processing Facility in Brevard County, FL. The ground controls remained at the SSPF for the entire mission duration. Scheduled operations began shortly after science loading and continued through launch, ascent, on-orbit, descent and ending upon return to Earth and the SSPF for deintegration. Space Tango’s Powered Ascent Utility Locker (PAUL) allows for CubeLabs to be fully powered while they are installed in the SpaceX Cargo Dragon vehicle. The PAUL provides the optimal culture environment and capabilities for the entire time the science remains within the CubeLab, including while on the launch pad and after splashdown. Flight and Ground systems for each mission were run synchronously with scheduled operations initiated at the exact same time.

The CubeLab culture environment was maintained at 37°C and 5-7% CO2, culture media and RNA-lysis buffer were maintained at 2-8°C providing optimal culture conditions. Media exchanges were performed every 7 days using microfluidic routing and pumping. Imaging of nanobioreactors was performed every day, for 30 days, beginning after installation on the International Space Station. SpaceX missions CRS-24, CRS-25, and CRS-27 on-orbit mission durations were 30 days, and the SpaceX CRS-26 on-orbit mission duration was 45 days. On the 30th day of on-orbit operations, cells were sampled from the nanobioreactors to the lysis buffer-containing bags using microfluidic routing and pumping of culture media. Lysis buffer was comprised of RLT Buffer (Qiagen) and 10% beta-mercaptoethanol. Media containing cells were flushed at a ratio of 1:3.5, media to lysis buffer.

CubeLab environmental telemetry (culture environment temperature, temperature of stored fluids, CubeLab CO2, CubeLab humidity) was recorded during powered operations. Telemetry data was available in near real-time through Space Tango’s customer dashboard.

### Imaging operations and post-processing (flight and ground, all missions)

Brightfield, green fluorescence, and red fluorescence imaging of each nanobioreactor were captured using Space Tango’s three-axis microscope system. A minimum of twelve imaging location presets were used for flight nanobioreactors and a minimum of five presets for ground nanobioreactors. Images were captured in monochrome and uploaded to Space Tango’s customer media portal for near real-time viewing during on-orbit operations.

Image post-processing was performed by Space Tango using ImageJ. A custom macro script was used to perform histogram adjustment, z-projection of z-slices, application of green and red LUTs to monochrome images, image composites, and brightfield, green and red image overlays. These post-processed image sets were uploaded to Space Tango’s customer media portal alongside the original monochrome images.

### Volocity Fluorescence Quantification

Still images taken during mission were analyzed using Volocity software licensed to the Microscopy Core at UC San Diego. Image Z-stack max projections were rendered in Image J. The max projections were rendered in grey-scale and exported as BMP files for analysis. Fluorescent signal was measured individually for mCherry and mVenus. Signal output was based on total fluorescing surface area (µm^2^) and plotted in GraphPad Prism.

### Colony Formation and Replating Assay

In vitro experiments were performed as previously described^48^. 10,000 of ABM cells were collected, resuspended in fresh media, and plated in methylcellulose (MC) (Stem Cell Technologies: MethoCult™ H4335 Catalog # 04330) in triplicates. After two weeks primary colonies (more than 40 cells) were counted and some individual multilineage colonies were plucked, resuspended and replated in fresh MC. Secondary colonies were counted after another 14 days. Data analysis: The absolute number of colonies was presented as the mean for triplicate conditions. Error bars indicate the SE. In some graphs, results are presented as summarized data (means) for four samples, and error bars indicate the SD. Statistical analysis included Student’s t-test and one-way Anova including All Pairwise Multiple Comparison Procedures (Holm-Sidak method).

### Sample Processing for Whole-Genome Sequencing

Recovered cells were pelleted and snap frozen at -80°C prior to DNA extraction. Pellets were processed using Qiagen DNeasy Blood and Tissue Kit according to manufacturers’ protocol (Qiagen, catalog 69504). DNA was eluted in Qiagen Buffer AE and frozen at -20°C. Samples were sequenced at 90x coverage for whole genome sequencing (NovoGene Corporation).

### Sample Processing for Whole-Transcriptome Sequencing

An aliquot of recovered cells was lysed in RLT buffer (Qiagen) and 10% beta-mercaptoethanol. Lysed samples were processed using Qiagen RNeasy Micro Kit according to manufacturers’ protocol (Qiagen, catalog 74004). RNA was eluted in RNase-free water and frozen at -80°C until RNA could be assessed for quality. Samples with RNA integrity number (RIN) >7 were further processed for RNA-sequencing (The Scripps Research Institute Next Generation Sequencing Core). RNA-Seq was performed on Illumina’s NextSeq 500 sequencer with 150bp paired-end reads.

### APOBEC3C lentiviral vectors and transduction

Lentiviral human wild-type APOBEC3C (pCDH-EF1-MCS-T2A-copGFP) was used as previously described^6^. All lentiviruses were tested by transduction of 293T cells and transduction efficiency was assessed by qRT-PCR. The cells were cultured for 72 hours in 96-well plates (2X105-5X105 cells per well) containing StemPro (Stem Cell Technologies) media supplemented with human IL-6, stem cell factor (SCF), Thrombopoietin (Tpo) and FLT-3 (all from R&D Systems)^36, 48-50^. The transduced cells were collected into RLT for RNA extraction or pelleted for DNA extraction.

### Whole-genome sequencing

Extracted DNA from these experiments was sent to Novogene (Sacramento, CA) for whole genome sequencing. DNA sequencing library preparation was performed using the NEBNext® DNA Library Prep Kit (New England Biolabs) following the manufacturer’s recommendations. Qualified libraries were paired and sequenced on an Illumina NovaSeq6000 platform using 150 PE sequencing chemistry to 90x coverage according to effective concentration and data volume.

### Identification of somatic mutations from whole genome bulk sequencing

Raw sequence data were downloaded to the Triton Shared Compute Cluster (TSCC) from ftp server link shared by Novogene (Sacramento, CA). All the post-sequencing analysis was performed within TSCC at UC San Diego. A schematic of the somatic mutations calling process is described in Supplementary fig S2. This methodology for identification of somatic mutations from bulk sequencing data follows established approaches from large genomics consortia (1). Briefly, quality assurance of the raw FASTQ files were evaluated using FastQC (Version 0.12.0) and Mosdepth (Version 0.3.4) (2, 3). Raw sequence reads were aligned to the human reference genome GRCh38.d1.vd1. The aligned reads were marked duplicated using MarkDuplicates (Picard) from GATK (Version 4.0.11.0) (4). An ensemble variant calling pipeline (EVC) was used to identify single nucleotide variants (SNV) and short insertions and deletions (indels). EVC implements the SNV and indel variant calling from four variant callers (Mutect2, Strelka2, Varscan2, and MuSE2) and only mutations that are identified by any two variant callers were considered as bona fide mutations (4-7).

### Telomere length estimates from whole genome bulk sequencing

A schematic of the Telomere length estimation process is described in Figs. 2a and S2a. The mark duplicated reads were used for telomere length estimation using Telomerecat (8) (Telomere Computational Analysis Tool). The ‘bam2length’ parameters were used with simulation sets to 100 and output options set to details parameters.

### Mitochondrial DNA copy number estimation from whole genome bulk sequencing

A schematic of the mitochondrial DNA copy number average process is described in Fig. S2. FastMitoCalc (9) tool was employed to determine the average mitochondrial DNA copy number by following the parameter described in the tools methods sections.

### Analysis of mutational profile

Analysis of mutational profiles was performed using our previously established methodology with the SigProfiler suite of tools used for summarization, simulation, visualization, and assignment of mutational profiles. Briefly, mutational matrixes for SBS, DBS and Indels were generated with SigProfilerMatrixGenerator (Version 1.2.16) (10). Plotting of each mutational profile was done with SigProfilerPlotting (Version 1.3.13).

### Code availability

Somatic mutations in whole-genome sequencing data were identified using our ensemble variant calling pipeline, which is freely available under the permissive 2-clause BSD license at: https://github.com/AlexandrovLab/EnsembleVariantCallingPipeline. All other computational tools utilized in this publication have been mentioned in the methodology section and can be accessed through their respective publications.

### RNA-Seq Analyses

RNA-Seq (2 x150bp) paired-end reads were generated and analyzed as described previously^25^. Briefly, following quality control with FastQC^51^ demultiplexed fastq files were aligned to the human genome, hg38, with STAR (v2.5.3a)^52^ with gene/transcript abundance quantified by RSEM (v1.3.0)^53^ GENCODE annotation (gencode.v43.annotation.gtf). Using functions from the R Bioconductor package edgeR^54^ and limma^55^ gene and isoform levels were separately aggregated for all samples, filtered for low expressing genes (counts per million < 1 in 50% of samples), Trimmed Mean of the M-values (TMM) normalized, and converted to log-transformed counts per million (lcpm) values for plotting. Enrichment analysis was performed with the R Bioconductor package fgsea^56^ and pathways from the Reactome database^57^.

### Quantification of Alternative Splicing

Bam files generated by STAR alignment to hg38 (described above) were used as input into the rMATS software (v4.1.0)^58^ to determine significant differential splicing events. For the event types Alterative 3’ Splice (A3’SS), Alternative 5’ Splice Site (A5’SS), Retained Intron (RI), and Skipped Exon (SE), events with FDR < 0.5 and |PSI| > 0.1 using both junction counts and exon counts were considered significant.

### RNA Editing

RNA editing events within each sample were determined as described previously^25^. ADAR-driven events were the focus of this analysis, so positive strand A to G and negative strand T to C variants were retained for quantification of total edits within each sample group.

### Repeat Element Analysis

Bam files generated by STAR alignment (described above), along with the repeatmasker.hg38.gtf file (downloaded from the UCSC genome browser), were used as input into the “featureCounts” function from the Subread package (https://subread.sourceforge.net/) to generate counts of repeat elements. As with the gene expression data, counts were aggregated for all samples, filtered, normalized, and converted lcpm. Repeat elements were also annotated using the “hg38.fa.out” file (downloaded from https://www.repeatmasker.org/species/hg.html). Differential expression of repeat elements was determined using limma^55^ and limma-voom^59^ with the model design (∼0 + flight_status).

### Mutation Annotation and Visualization

Somatic mutations from whole genome sequencing data were later converted to MAF using vcf2maf (Cyriac Kandoth. mskcc/vcf2maf: vcf2maf v1.6.19. (2020). doi:10.5281/zenodo.593251) utility. Oncoplot were generated using Maftools^60^ R Bioconductor package.

## Acknowledgments

We would like to acknowledge Ara Lidstrom and Michelle Ghani, Executive Director of the Sanford Stem Cell Institute for administrative support.

UCSD Microscopy Core, Grant NINDS P30NS047101

## Funding

We would like to thank our funding agencies for their vital support, including grants from:

NASA (NRA NNJ13ZBG001N) to Space Tango and UC San Diego ISSCOR CHMJ

NIH/NCI (R01CA205944) to CHMJ

NIH/NIDDK (R01DK114468-01) to CHMJ

LLS Blood Cancer Discoveries

Sanford Stem Cell Institute

Koman Family Foundation

JM Foundation

Moores Family Foundation.

## Author contributions

Conceptualization: CHMJ, LL, JP, LB

Methodology: CHMJ, JP, JI, LL, LB, SN, EK, AR, DM, PG, SG, JJ, TN, TM, AK, JS, MPS, TC, ARM, SRM, TW, LB, LA

Investigation: JP, LB, JI, LL, TN, DM, SN, TW, WCB

Visualization: JP, TN, DM, SN, TW

Funding acquisition: CHMJ, JS, ARM, SRM, TC, KM

Project administration: CHMJ, TC

Supervision: CHMJ, TC

Writing - original draft: CHMJ, KM, JP

Writing - review & editing: CHMJ, KM, JP, JI, LB, LL, SN, TW, LA, TN, DM, JJ, TC, AK, IvdW

## Data and materials availability

Documentation for the computational analyses will be made available at https://github.com/ucsd-ccbb/MPN_atlas_methods. Sequencing data sets will be available through dbGAP. Additional data are available from the corresponding author upon reasonable request.

## References

1. Rossi, D.J., Jamieson, C.H.M. & Weissman, I.L. Stems Cells and the Pathways to Aging and Cancer. Cell 132, 681–696 (2008).

2. López-Otin, C., Blasco, M.A., Partridge, L., Serrano, M. & Kroemer, G. Hallmarks of aging: An expanding universe. Cell 186, 243–278 (2023).

3. Chua, B.A., Van Der Werf, I., Jamieson, C. & Signer, R.A.J. Post-Transcriptional Regulation of Homeostatic, Stressed, and Malignant Stem Cells. Cell Stem Cell 26, 138–159 (2020).

4. Jamieson, C. Bad blood promotes tumour progression. Nature 549, 465–466 (2017).

5. Waclawiczek, A. et al. Combinatorial BCL2 Family Expression in Acute Myeloid Leukemia Stem Cells Predicts Clinical Response to Azacitidine/Venetoclax. Cancer Discovery 13, 1408–1427 (2023).

6. Jiang, Q. et al. Inflammation-driven deaminase deregulation fuels human pre-leukemia stem cell evolution. Cell Rep 34, 108670 (2021).

7. Solomon, O. et al. Global regulation of alternative splicing by adenosine deaminase acting on RNA (ADAR). RNA 19, 591–604 (2013).

8. Ota, H. et al. ADAR1 forms a complex with Dicer to promote microRNA processing and RNA-induced gene silencing. Cell 153, 575–589 (2013).

9. Lazzari, E. et al. Alu-dependent RNA editing of GLI1 promotes malignant regeneration in multiple myeloma. Nat Commun 8, 1922 (2017).

10. Jiang, Q. et al. Hyper-Editing of Cell-Cycle Regulatory and Tumor Suppressor RNA Promotes Malignant Progenitor Propagation. Cancer Cell 35, 81–94 e87 (2019).

11. Alexandrov, L.B. et al. The repertoire of mutational signatures in human cancer. Nature 578, 94–101 (2020).

12. Burns, M.B. et al. APOBEC3B is an enzymatic source of mutation in breast cancer. Nature 494, 366–370 (2013).

13. Freed, L.E., Langer, R., Martin, I., Pellis, N.R. & Vunjak-Novakovic, G. Tissue engineering of cartilage in space. Proceedings of the National Academy of Sciences 94, 13885–13890 (1997).

14. Pleguezuelos-Manzano, C. et al. Mutational signature in colorectal cancer caused by genotoxic pks+ E. coli. Nature 580, 269–273 (2020).

15. Poore, G.D. et al. Microbiome analyses of blood and tissues suggest cancer diagnostic approach. Nature 579, 56–574 (2020).

16. Steele, C.D. et al. Signatures of copy number alterations in human cancer. Nature 606, 984–991 (2022).

17. Alexandrov, L.B. et al. Mutational signatures associated with tobacco smoking in human cancer. Science 354, 618–622 (2016).

18. Alexandrov, L.B. et al. Clock-like mutational processes in human somatic cells. Nat Genet 47, 1402–1407 (2015).

19. Alexandrov, L.B., Nik-Zainal, S., Siu, H.C., Leung, S.Y. & Stratton, M.R. A mutational signature in gastric cancer suggests therapeutic strategies. Nat Commun 6, 8683 (2015).

20. Alexandrov, L.B. et al. Signatures of mutational processes in human cancer. Nature 500, 415–421 (2013).

21. Alexandrov, L.B. et al. The repertoire of mutational signatures in human cancer. Nature 578, 94–101 (2020).

22. Shiromoto, Y., Sakurai, M., Minakuchi, M., Ariyoshi, K. & Nishikura, K. ADAR1 RNA editing enzyme regulates R-loop formation and genome stability at telomeres in cancer cells. Nature Communications 12, 1654 (2021).

23. Blackburn, E.H. Cancer Interception. Cancer Prevention Research 4, 787–792 (2011).

24. Crews, L.A. et al. Reversal of malignant ADAR1 splice isoform switching with Rebecsinib. Cell Stem Cell (2023).

25. van der Werf, I. et al. Detection and targeting of splicing deregulation in pediatric acute myeloid leukemia stem cells. Cell Reports Medicine (2023).

26. Garrett-Bakelman, F.E. et al. The NASA Twins Study: A multidimensional analysis of a year-long human spaceflight. Science 364 (2019).

27. Wnorowski, A. et al. Effects of Spaceflight on Human Induced Pluripotent Stem Cell-Derived Cardiomyocyte Structure and Function. Stem Cell Reports 13, 960–969 (2019).

28. Oh, J., Lee, Y.D. & Wagers, A.J. Stem cell aging: mechanisms, regulators and therapeutic opportunities. Nature Medicine 20, 870–880 (2014).

29. Scott, J.M., Stoudemire, J., Dolan, L. & Downs, M. Leveraging Spaceflight to Advance Cardiovascular Research on Earth. Circulation Research 130, 942–957 (2022).

30. Pineda, G. et al. Tracking of Normal and Malignant Progenitor Cell Cycle Transit in a Defined Niche. Scientific reports 6, 23885 (2016).

31. Lemons, D.E. & Downey, J.A. in The Physiological Basis of Rehabilitation Medicine (Second Edition). (eds. J.A. Downey, S.J. Myers, E.G. Gonzalez & J.S. Lieberman) 365–391 (Butterworth-Heinemann, 1994).

32. Gilley, D., Herbert, B.-S., Huda, N., Tanaka, H. & Reed, T. Factors impacting human telomere homeostasis and age-related disease. Mechanisms of Ageing and Development 129, 27–34 (2008).

33. Luxton, J.J. et al. Telomere Length Dynamics and DNA Damage Responses Associated with Long-Duration Spaceflight. Cell Reports 33 (2020).

34. Alexandrov, L.B. et al. Signatures of mutational processes in human cancer. Nature 500, 415–421 (2013).

35. Gorbunova, V. et al. The role of retrotransposable elements in ageing and age-associated diseases. Nature 596, 43–53 (2021).

36. Zipeto, Maria A. et al. ADAR1 Activation Drives Leukemia Stem Cell Self-Renewal by Impairing Let-7 Biogenesis. Cell Stem Cell 19, 177–191 (2016).

37. Gambino, F. et al. IL1RAPL1 controls inhibitory networks during cerebellar development in mice. European Journal of Neuroscience 30, 1476–1486 (2009).

38. Malloy, C., Ahern, M., Lin, L. & Hoffman, D.A. Neuronal Roles of the Multifunctional Protein Dipeptidyl Peptidase-like 6 (DPP6). International Journal of Molecular Sciences 23, 9184 (2022).

39. Montani, C. et al. The X-Linked Intellectual Disability Protein IL1RAPL1 Regulates Dendrite Complexity. The Journal of Neuroscience 37, 6606–6627 (2017).

40. Yokoyama, K. et al. NYAP: a phosphoprotein family that links PI3K to WAVE1 signalling in neurons. The EMBO Journal 30, 4739–4754 (2011).

41. Gao, X. et al. Nociceptive nerves regulate haematopoietic stem cell mobilization. Nature 589, 591–596 (2021).

42. Scott EC G.E., Masood A, Chuang NT, Vertino PM, Devine SE A hot L1 retrotransposon evades somatic repression and initiates human colorectal cancer. Genome Res 26, 745–755 (2016).

43. Schumacher, A.J., Haché, G., MacDuff, D.A., Brown, W.L. & Harris, R.S. The DNA Deaminase Activity of Human APOBEC3G Is Required for Ty1, MusD, and Human Immunodeficiency Virus Type 1 Restriction. Journal of Virology 82, 2652–2660 (2008).

44. Ramini, D. et al. Replicative Senescence-Associated LINE1 Methylation and LINE1-Alu Expression Levels in Human Endothelial Cells. Cells 11, 3799 (2022).

45. Bisserier, M. et al. Cell‐Free Mitochondrial DNA as a Potential Biomarker for Astronauts’ Health. Journal of the American Heart Association 10, e022055 (2021).

46. Nguyen, H.P., Tran, P.H., Kim, K.-S. & Yang, S.-G. The effects of real and simulated microgravity on cellular mitochondrial function. npj Microgravity 7, 44 (2021).

47. Hematology, A.S.o. (ed. K. Ladel L, Oliver I, Jamieson C. ) (Blood 2022;1277, 2022).

48. Goff, Daniel J. et al. A Pan-BCL2 Inhibitor Renders Bone-Marrow-Resident Human Leukemia Stem Cells Sensitive to Tyrosine Kinase Inhibition. Cell Stem Cell 12, 316–328 (2013).

49. Abrahamsson, A.E. et al. Glycogen synthase kinase 3β missplicing contributes to leukemia stem cell generation. Proceedings of the National Academy of Sciences 106, 3925–3929 (2009).

50. Jiang, Q. et al. ADAR1 promotes malignant progenitor reprogramming in chronic myeloid leukemia. Proceedings of the National Academy of Sciences 110, 1041–1046 (2013).

51. Wingett, S. & Andrews, S. FastQ Screen: A tool for multi-genome mapping and quality control [version 2; peer review: 4 approved]. F1000Research 7 (2018).

52. Dobin, A. et al. STAR: ultrafast universal RNA-seq aligner. Bioinformatics 29, 15–21 (2012).

53. Li, B. & Dewey, C.N. RSEM: accurate transcript quantification from RNA-Seq data with or without a reference genome. BMC Bioinformatics 12, 323 (2011).

54. Robinson, M.D., McCarthy, D.J. & Smyth, G.K. edgeR: a Bioconductor package for differential expression analysis of digital gene expression data. Bioinformatics 26, 139–140 (2009).

55. Ritchie, M.E. et al. limma powers differential expression analyses for RNA-sequencing and microarray studies. Nucleic Acids Research 43, e47–e47 (2015).

56. Korotkevich, G. et al. Fast gene set enrichment analysis. bioRxiv, 060012 (2021).

57. Gillespie, M. et al. The reactome pathway knowledgebase 2022. Nucleic Acids Research 50, D687–D692 (2021).

58. Shen, S. et al. rMATS: Robust and flexible detection of differential alternative splicing from replicate RNA-Seq data. Proceedings of the National Academy of Sciences 111, E5593–E5601 (2014).

59. Law, C.W., Chen, Y., Shi, W. & Smyth, G.K. voom: precision weights unlock linear model analysis tools for RNA-seq read counts. Genome Biology 15, R29 (2014).

60. Mayakonda, A., Lin, D.-C., Assenov, Y., Plass, C. & Koeffler, H.P. Maftools: efficient and comprehensive analysis of somatic variants in cancer. Genome research 28, 1747–1756 (2018).

