## Supplemental for "Accelerated Hematopoietic Stem Cell Aging in Space"

**List of Supplementary Materials:**

Figs. S1 to S4

Supplemental Figure 1

A

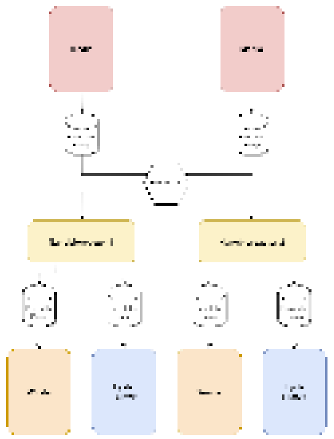

B

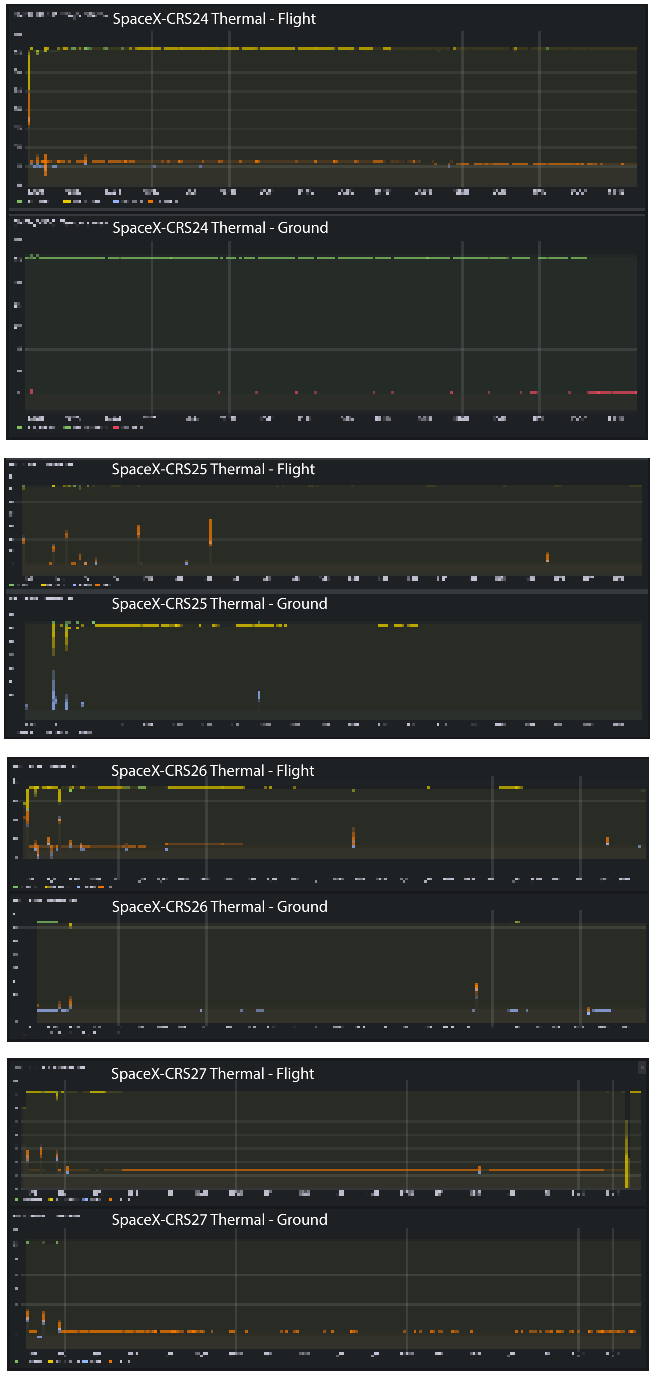


C

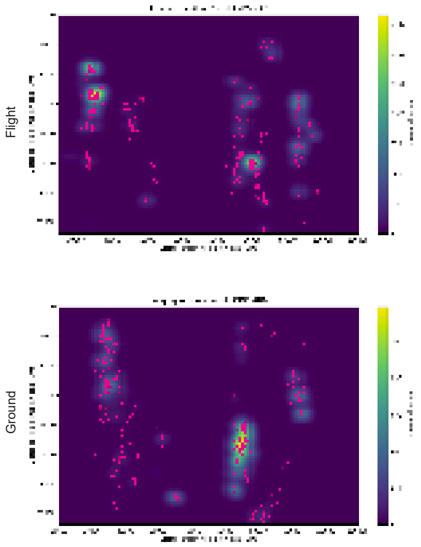


D

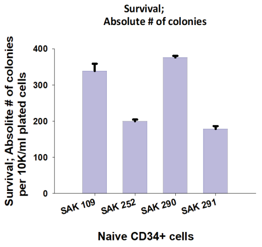


E

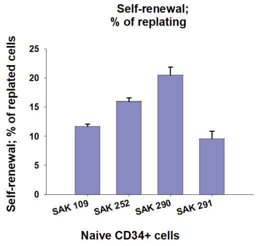


F

| Sample | Condition | %Viability |
|---|---|---|
| **SpX-CRS24_SAK109** | Flight | 69% |
| **SpX-CRS24_SAK109** | Ground | 51% |
| **SpX-CRS24_SAK066** | Flight | 70% |
| **SpX-CRS24_SAK072** | Ground | 83% |
| **SpX-CRS26_SAK252** | Flight | 32% |
| **SpX-CRS26_SAK252** | Ground | 30% |
| **SpX-CRS27_SAK290** | Flight | 16% |
| **SpX-CRS27_SAK290** | Ground | 48& |
| **SpX-CRS27_SAK291** | Flight | 27% |
| **SpX-CRS27_SAK291** | Ground | 21% |

G

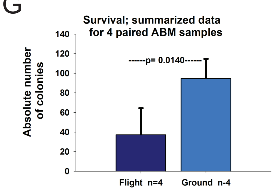


H

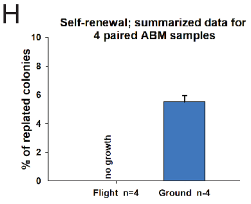

**Figure S1. Pre-, during, and post-flight mission operations**

(**A**) Space Tango CubeLab hardware fluidics schematic used for all four missions.

(B) Plots of the temperatures of biological incubator and media for on-orbit and terrestrial experiments over the entirety of respective experiment duration, ordered by flight.

(C) Space Tango preset summary of flight-combined and ground-combined locations that imaging occurred daily over the course of four missions.

(D-E) Four individual aged bone marrow (ABM) sample naïve CD34+ cells were directly subjected to survival and self-renewal assays to evaluate the maximum level of their clonogenic ability. (D) Represents the results of performed survival assay (primary colonies). The graph demonstrates the absolute total number of colonies. (E) Represents results of self-renewal assays (secondary colonies). The graph depicts the % of replated cells. Each bar represents Mean+/- SE for triplicate conditions.

(F) Percent viability of paired (N=4) and unpaired (N=2) samples from four SpX-CRS missions upon return. Viability was measured with TC20 Automated Cell Counter using trypan blue (BIO-RAD).

(G-H) Summarized data of four individual paired ABM-derived cells (SAK-109, SAK-252, SAK-290, and SAK-291) that were cultured in parallel in flight and ground nanobioreactors for 42 days. Upon return, cells were subjected to clonogenic assays, as described in Fig. 1A and Methods. Data (means) for summarized four flight or ground samples respectively are presented. Error bars indicate the SD. Statistical analysis included Student's t-test and one-way ANOVA, including All Pairwise Multiple Comparison Procedures (Holm-Sidak method).

Supplemental Figure 2

A

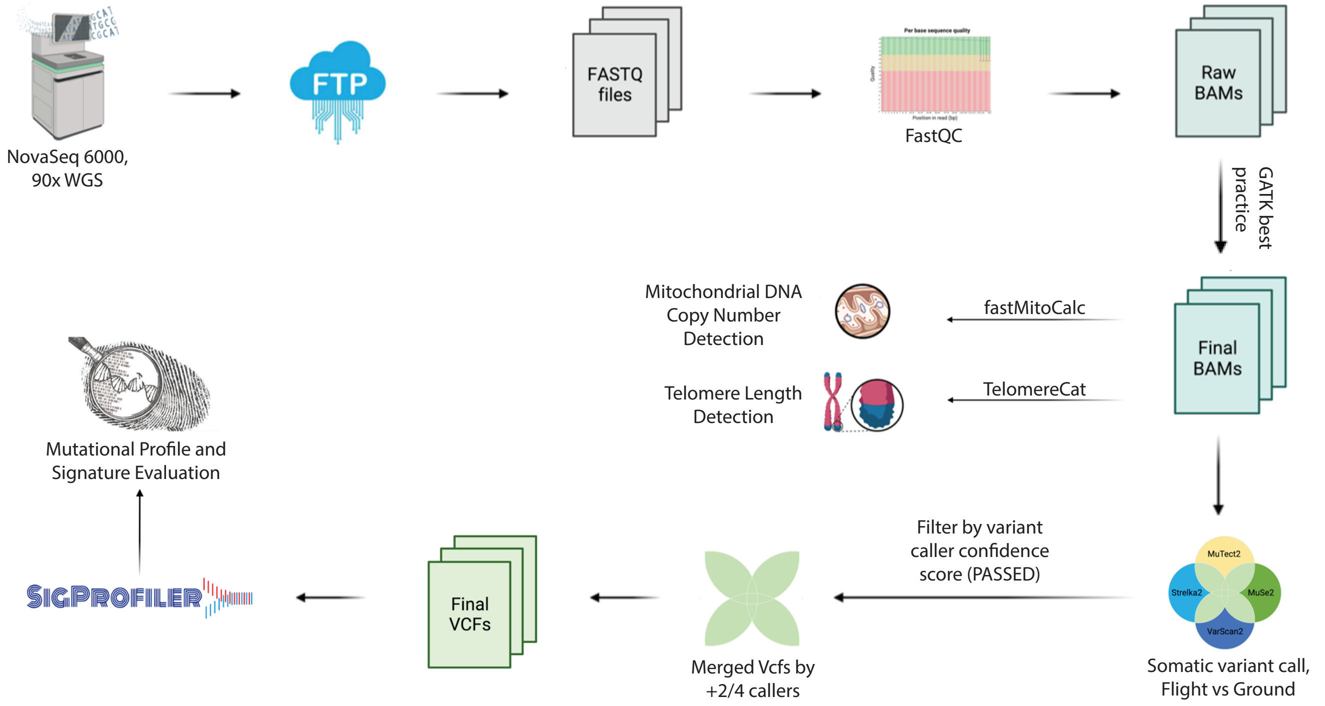


B

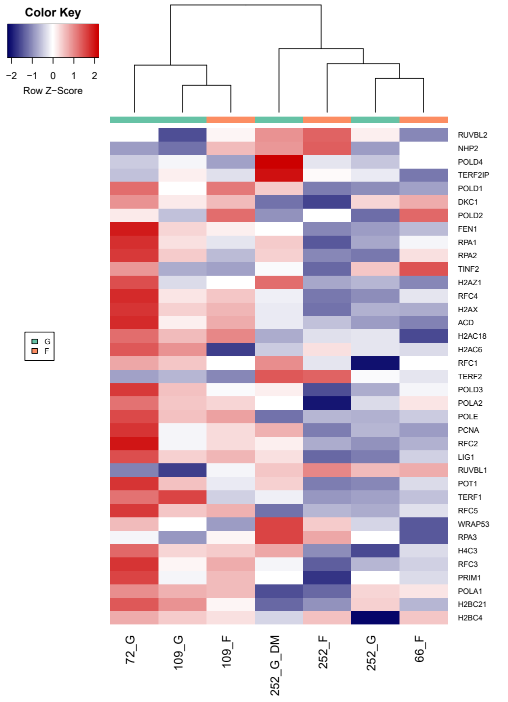


C

Mitochondrial Translation

| Gene | logFC |
|---|---|
| H2BC21 | -1.55 |
| TOP3A | -0.76 |
| CLASP1 | -0.59 |
| DYNLL2 | 0.55 |
| RMI2 | 0.88 |

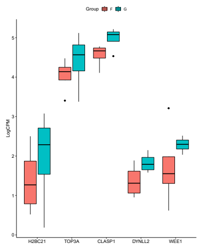

D

Interferon alpha/beta Signaling

| Gene | logFC |
|---|---|
| IRF7 | -1.42 |
| SP100 | -0.69 |
| RSAD2 | -3.18 |
| ADAR | -0.73 |
| JAK2 | -0.53 |

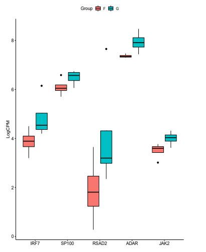

E

Cell Cycle Checkpoints

| Gene | logFC |
|---|---|
| MRPL12 | 0.86 |
| MRPL54 | 0.61 |
| MRPS34 | 0.49 |
| MRPS27 | 0.57 |
| MRPS17 | 0.59 |

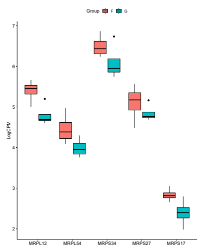

**Figure S2. Mitochondrial and telomere changes post flight**

(**A**) Schematic of whole-genome sequence data analysis. Raw FASTQ files were downloaded within Triton shared computational cluster environment. Sequencing reads quality was checked with FASTQC. BWA MEM was used to align the raw reads with GRCh38.d1.vd1 reference sequences. Following GATK best practice, four variant callers (Mutect2, VarScan2, Strelka2, and MuSE2) were employed in matching normal mode to call somatic mutations. Only mutations that are identified by any two variant callers were considered bona fide mutations. The final set of somatic mutations was analyzed by the SigProfiler suite of tools. Final binary alignment map (BAM) files were also analyzed via fastMitoCalc and TelomereCat tools to detect mitochondrial DNA copy number and telomere length estimation, respectively.

(B) Heatmap visualizing expression levels of telomerase-related genes, based on the telomerase Reactome pathway, in hematopoietic stem and progenitor cells (HSPCs) derived from ground and flight samples upon return.

(C-E) GSEA results for the top pathways by adjusted p-value with a positive and negative normalized enrichment score (NES), which include cell cycle checkpoints, mitochondrial translation and interferon alpha/beta signaling.

Supplemental Figure 3

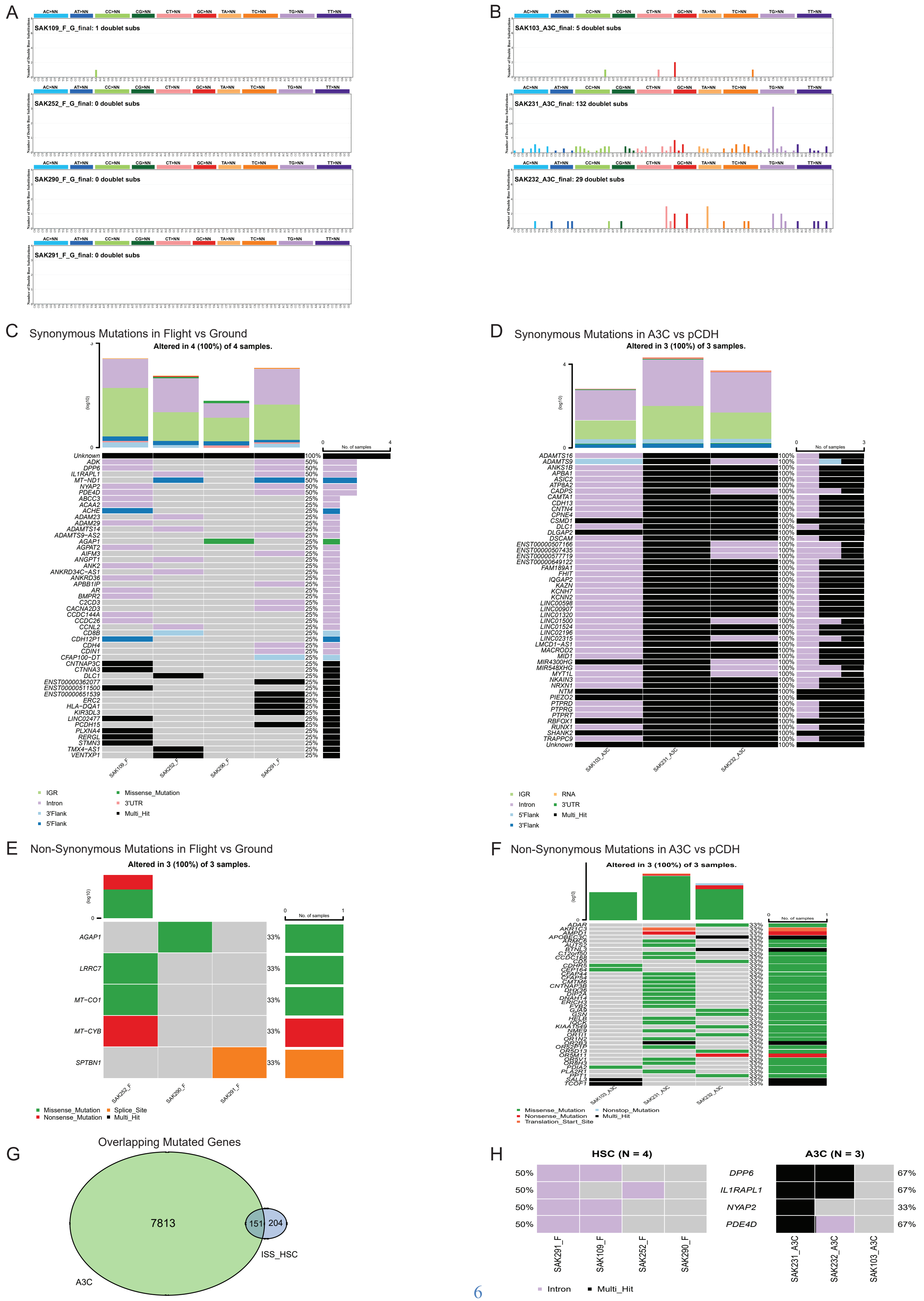

**Figure S3. Double base substitution mutation profile**

(**A**) Patterns of doublet base substitution (DBS) for samples from hematopoietic stem and progenitor cells (HSPCs) are shown using the DBS-78 classification scheme on the x-axis. The Y-axis are scaled differently in each plot to optimally show each mutational pattern with the y-axis reflecting the number of mutations for the respective mutational scheme.

(B) Patterns of doublet base substitution (DBS) for samples from APOBEC3C (A3C) expressed cells are shown using the SBS96 classification scheme on the x-axis. The Y-axis are scaled differently in each plot to optimally show each mutational pattern with the y-axis reflecting the number of mutations for the respective mutational scheme.

(C) Oncoplot displaying all the somatically mutated genes within HSPCs upon return. The data from returned flight samples is normalized to returned ground data. Genes are ordered by their mutation frequency, and samples are ordered numerically (bottom). The top bar plot shows log10 transformed mutational burden.

(D) Oncoplot displaying all the somatically mutated genes within APOBEC3C (A3C) overexpressing HSPCs. Data from A3C overexpressing cells is normalized to backbone, PCDH, transduced cells. Genes are ordered by their mutation frequency, and samples are ordered numerically (bottom). The top bar plot shows log10 transformed mutational burden.

(E) Oncoplot displaying non-synonymously mutated genes within HSPCs upon return. The data from returned flight samples is normalized to returned ground data. Genes are ordered by their mutation frequency, and samples are ordered numerically (bottom). The top bar plot shows log10 transformed mutational burden.

(F) Oncoplot displaying non-synonymously mutated genes within APOBEC3C (A3C) overexpressing HSPCs. Data from A3C overexpressing cells is normalized to backbone, PCDH, transduced cells. Genes are ordered by their mutation frequency, and samples are ordered numerically (bottom). The top bar plot shows log10 transformed mutational burden.

(G) All overlapping mutated genes between HSPCs found in flight samples upon return, and APOBEC3C (A3C) cells.

(H) Differentially mutated genes between HSPC and APOBEC3C displayed as a forest plot. The top four mutated genes shared between the HSPC and APOBEC3C groups are shown.

Supplemental Figure 4

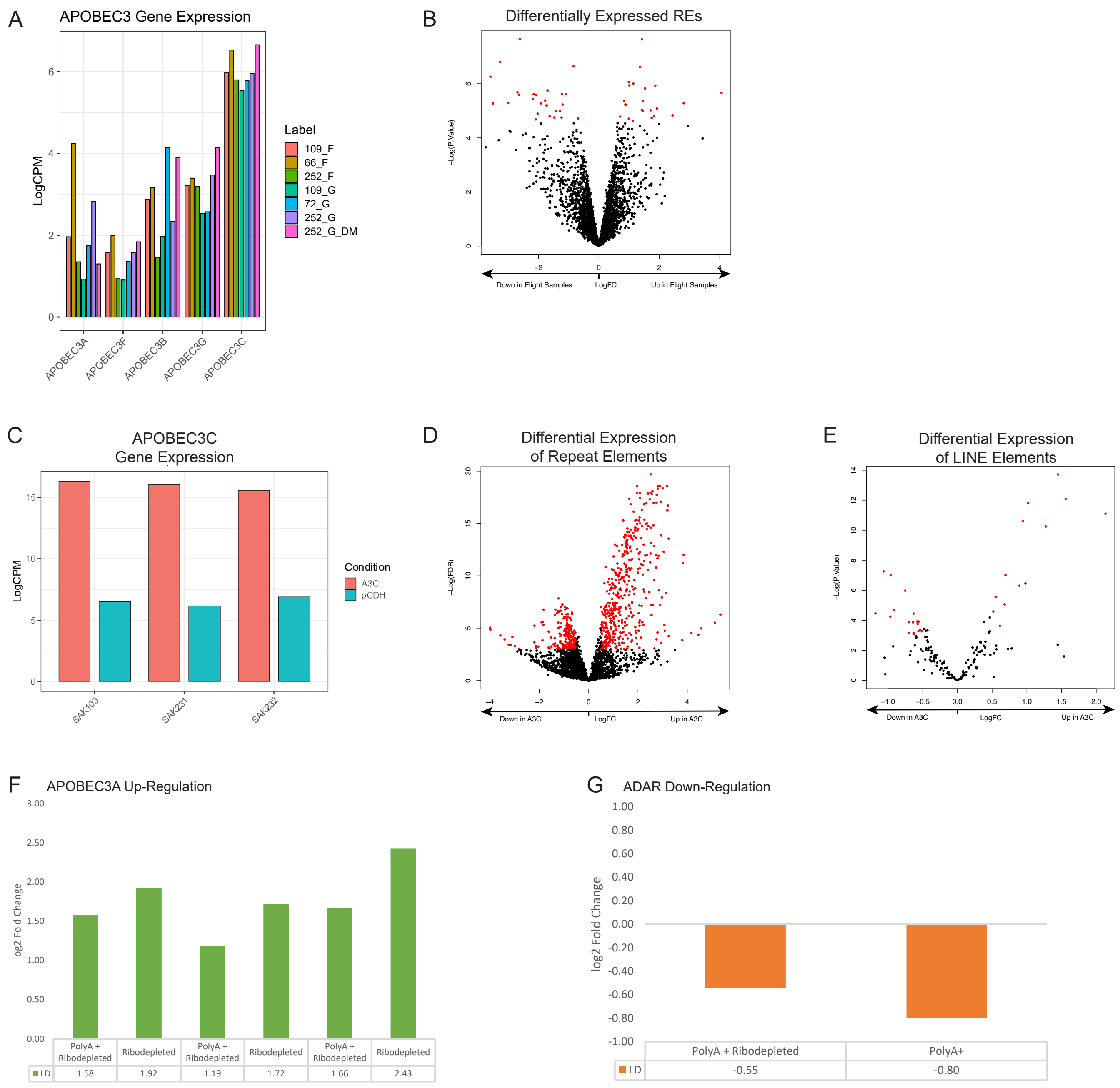

**Figure S4. Correlated gene expression changes**
(**A**) Barplot of all expressed APOBEC3 genes in individual flight and ground nanobioreactors sampled upon return from mission. Sample 252_G_DM was sampled during mission.
(B) Volcano plot visualizing expression of repetitive elements (RE) in hematopoietic stem and progenitor cells (HSPCs) derived from flight samples in comparison to ground samples upon return. Y-axis represents the negative log of the p-value (-Log P Value) and the x-axis represents the log fold change (LogFC). Red dots represent repetitive elements with p-value < 0.01 used in (Fig. 4F).
(C) Barplot of APOBEC3C expression levels in APOBEC3C (A3C) overexpressing and backbone (pCDH) transduced aged bone marrow (ABM) samples.
(D) Volcano plot visualizing expression levels of repeat elements (repetitive elements) in APOBEC3C (A3C) overexpressing cells in comparison to backbone (pCDH) transduced cells. Red dots represent elements with adjusted p-value < 0.05 and logFC > 0.5 or < 0.05.
(E) Volcano plot visualizing expression levels of Long Interspersed Nuclear Elements (LINEs) in APOBEC3C (A3C) overexpressing cells in comparison to backbone (pCDH) transduced cells. Red dots represent elements with adjusted p-value < 0.05 and logFC > 0.5 or < 0.05.
(F) Barplot of APOBEC3A expression levels in lymphocyte-depleted cells as time in space increases.
(G) Barplot of ADAR1 expression levels in lymphocyte-depleted cells as time in space increases. Note, in this analysis, expression levels of ADAR1 splice isoforms ADAR1p150 and ADAR1 p110 was not determined.
